# Soil respiration variation along an altitudinal gradient in Italian Alps: Disentangling forest structure and temperature effects

**DOI:** 10.1101/2021.02.17.431600

**Authors:** Aysan Badraghi, Maurizio Ventura, Andrea Polo, Luigimaria Borruso, Leonardo Montagnani

## Abstract

To understand the main determinants of soil respiration (SR), we investigated the changes of soil respiration and soil physicochemical properties, including soil carbon (C) and nitrogen (N), root C and N, litter C and N, soil bulk densities and soil pH at five forest sites, along an elevation/temperature gradient (404 to 2101 m a.s.l) in Northern Italy, where confounding factors such as aspect and soil parent material are minimized, but an ample variation in forest structure and composition is present. Our result indicated that SR rates increased with temperature in all sites, and about 55% - 76% of SR was explained by temperature. Annual cumulative SR, ranging between 0.65 and 1.40 kg C m^-2^ yr^-1^, declined along the elevation gradient, while temperature sensitivity (Q10) of SR increased with elevation. However, a high SR rate (1.27 kg C m^-2^ yr^-1^) and low Q10 were recorded in the old conifer forest stand at 1731 m a.s.l., characterized by a complex structure and high productivity, introducing nonlinearity in the relations with elevation and temperature. Reference SR at the temperature of 10°C (SR_ref_) was not related to elevation. A significant linear negative relationship was found for bulk density with elevation. On the contrary, soil C, soil N, root C, root N, pH and litter mass were better fitted by nonlinear relations with elevation. However, it was not possible to confirm a significant correlation of SR with these parameters once the effect of temperature has been removed (SR_ref_). These results show how the main factor affecting SR in forest ecosystems along this Alpine elevation gradient is temperature, but its regulating role can be strongly influenced by site biological characteristics, particularly vegetation type and structure. This study also confirms that high elevation sites are rich in C stored in the soil and also more sensitive to climate change, being prone to high carbon losses as CO_2_. Conversely, forest ecosystems with a complex structure, with high SR_ref_ and moderate Q10, can be more resilient.

## Introduction

Soil respiration (SR) is the largest biological carbon (C) flux after photosynthesis in terrestrial ecosystems [1]. It largely determines the C balance between the terrestrial biosphere and the atmosphere [2,3,4] and assumes a decisive role in the carbon cycle and terrestrial carbon sink capacity. The soil is the largest C pool in the terrestrial biosphere and has been increasingly recognized to play a crucial role in mitigating global warming resulting from climate change [5, 6, 7]. Small changes in soil CO2 efflux or soil organic C stocks could severely impact the global C cycle [8]. In this regard, SR is one of the fluxes that have received more attention by research for a longer time. The study by Janssens et al. [9] evidenced the relevant role of forest productivity in the determination of SR. Since then, other studies have investigated and quantified the impact of productivity on SR modeling [10, 11]. Apart from productivity, SR is influenced by different abiotic and biotic factors such as soil temperature, moisture, and microbial community, introducing a considerable uncertainty in SR estimates [12,13,14]. Among these factors, the temperature has been the most often studied factor affecting respiratory processes [15,16]. Predicting the SR response to increasing temperature (temperature sensitivity of SR) has been one of the main objectives of research for years; therefore different equations relating soil CO2 efflux with temperature [17,18,19] or with a combination of temperature and soil humidity have been developed [20]. Nevertheless, the Q10 function [21] using the Q10 parameter to describe the temperature sensitivity of SR is one of the most widely used models, still mainly employed to quantify the CO2 efflux from the soil in Earth system models.

On the other hand, the elevation is a key driver of climate properties. It plays an essential role in the soil organic matter distribution and may dampen the effects of climate change [14,22,23,24, 25]. In general, temperature declines with elevation, thus elevation gradient has been used to assess soil respiration response to temperature in several studies [12,26,27,28]. These studies indicate that CO2 exchange between soil and atmosphere varies along climatic gradients and that temperature sensitivity (Q10) of SR increases with elevation. They also found a positive relationship between soil organic matter (SOM) and elevation and reported that global soil organic C stock at high elevation is more sensitive to climate change and is predicted to decrease in a warming climate [14,23,29,30,31,32]. However, several researchers have reported opposite trends and found lower SOM and higher SR at a higher elevation [30,33,34]. This variability may be partially due to confounding factors affecting SR other than temperature. Besides elevation, mountain landscapes are, in fact, characterized by substantial changes of other site parameters such as slope and aspect, which can affect microclimatic conditions and, therefore soil C dynamics [35]. Furthermore, due to the heterogeneity of geological substrates, soils of mountain regions are highly diverse over short spatial scales and this can generate marked contrasts in soil biogeochemical functions [36]. Different results have also been found when SR is related to soil organic carbon [26,37].

Besides, there is evidence that diverse plant biome types can influence SR rate differently. Therefore, the various plant communities can affect differently, microclimate, soil and litter composition, and root distribution, therefore affecting soil respiration rate [18, 26,38,39,40]. However, within the same plant biome, there is a high spatial heterogeneity of SR. Some authors found a possible linkage between the topography, plant community structure (e.g., forest type and speed of regeneration), and SR within the same forest ecosystems [18,38,39,40]. Further, forest management can also play a crucial role in SR [41]. For instance, tree removal can directly influence soil respiration due to the removal itself (i.e., reduction of plant biomass) but even indirectly changing the soil’s physicochemical properties and micrometeorological conditions [42].

Currently, the temperature dependency of SR and SOC decomposition is a major interest regarding global climate change and the role of terrestrial ecosystems in regulating Earth’s climate [43,44]. Therefore, there is a need to better understand the interactions between temperature and soil CO2 efflux. The general goal of this study is to disentangle the possible multi-effects on SR of soil properties, temperature, SOM, and vegetation structure (tree height in particular) along a plant biome-elevation gradient. In particular, the existing differences in vegetation structure allowed us to investigate the extent to which these biological variables and the induced variation in microclimatology can alter the relation between elevation and SR.

Specifically, i), we tested the hypothesis that SR and SOM accumulation change linearly with elevation. We also hypothesized that the Q10 value increases linearly with elevation as well. Furthermore, ii) we analyzed which are the main factors affecting SR other than temperature. To better isolate the effect of temperature on SR, the study was conducted along an altitudinal/temperature gradient in Italian Alps, in conditions where confounding factors like slope, aspect, and soil parent material are minimized. The differences in vegetation structure allowed us to investigate to which extent these biological variables and the induced variation in microclimatology, can alter the relation between elevation and SR.

## Material and Methods

### Study areas

Five experimental sites were established between the top of the Rittner Horn mount and the city of Bolzano, Italy, on the southern side of the Alps (Fig 1a). The overall elevation gradient between the highest and the lowest site is 1700 m and the elevation separation between each site is approximately 420 ± 60 m. All sites are characterized by a soil evolved upon a glacial till laid on a porphyric bedrock and an SSE slope orientation. Annual precipitation ranges between 800 and 1000 mm.

**Fig 1.**
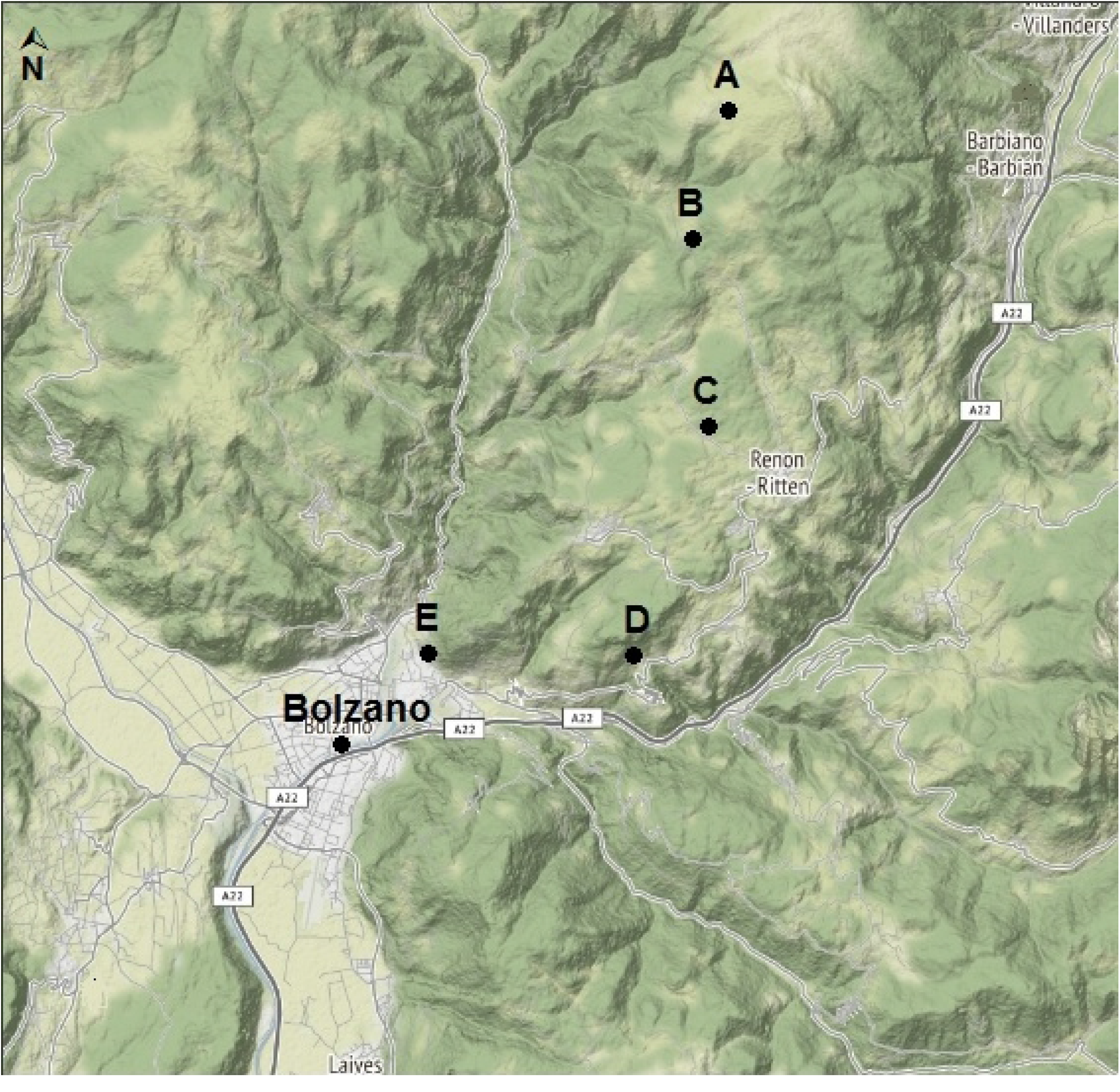
(a) Map showing the research site locations selected for this study (b) Scheme of the elevation and vegetation gradient present along the studied Alpine slope.

Site A was established in shrubland vegetation of Dwarfing Mountain pine (*Pinus mugo* Turra) near the summit of the Rittnern Horn/Corno del Renon mount (see the scheme in Fig 1b). Site B was established in a Norway spruce stand (*Picea abies* (L.) Karst.) at the long-term Fluxnet research station of Renon-Mittelgrünwald/Selva Verde (https://doi.org/10.18140/flx/1440173;). The site is characterized by an unvenaged distribution of tree diameters, approaching the structure of old-growth forest stands (45,46]. Site C was located near the location of Riggermoos (Oberbozen/Sopra Bolzano), in a low-density Scots pine (*Pinus sylvestris* L:) stand. Site D was established in a mixed stand of Sessile oak (*Quercus petrea* (Matt.) Liebl.) and Chestnut (*Castanea sativa* L:) and the presence of Scots pine near the village of Signat/Signato. Site E was located in a stand dominated by Downy oak (*Quercus pubescens* Willd.) and Flowering ash (*Fraxinus ornus* L.) on the hill slope just close to the city of Bolzano (Sankt Magdalena/Santa Maddalena). All sites except site A are managed as high forest mainly for wood harvesting. Site A is managed as natural vegetation with occasional harvesting only at forest margins to avoid expanding the pines in the adjacent pasture areas. Tree age was assessed in 2018 by tree ring count: it was found that site B had the oldest trees, slightly above 200 years, while other stands were in the range of 50-100 years. Tree height was assessed in 2020 with the TruePulse sensor (TruPulse 360 B laser range-finder, Laser Tech, Colorado, USA). Details on tree heights and the main characteristics of the research sites are reported in Table 1.

**Table 1.** General characterization of the study sites.

| Characteristics | Site A | Site B | Site C | Site D | Site E |
| --- | --- | --- | --- | --- | --- |
| Elevation<br>(m a.s.l) | 2101 | 1731 | 1354 | 865 | 404 |
| Mean annual<br>Temperature | 4 | 4 | 12 | 11 | 14 |
| (°C) |  |  |  |  |  |
| Land use | Shrubland | Forest | Forest | Forest | Forest |
| Mean dominant tree height (m) | 1.5 | 29 | 18 | 15 | 5 |
| Dominant species in the overstory | - Dwarfing Mountain pines (*Pinus mugo*) | - Norway spruce (*Picea abies*)<br>- Swiss stone pine (*Pinus cembra*)<br>- Larch (*Larix decidua*) | - Scots pine (*Pinus sylvestris*) | - Sessile oak (*Quercus petraea*)<br>- Scots Pine (*Pinus sylvestris*)<br>- Chestnut (*Castanea sativa*) | - Downy oak (*Quercus pubescens*)<br>- Flowering ash (*Fraxinus ornus*) |
| Main species in the understory | - Intervening grasses (*Festuca halleri*) | - Rusty leaved alprose (*Rhododendron ferrugineum*) | - Heather (*Erica carnea*) | Understory almost absent | - Smoke-bush (*Cotinus coggyria*)<br>- Succulent plants (*Opuntia humifusa*) |

### Soil respiration measurements

To quantify SR, ten iron collars (10 cm height, 20 cm diameter) were inserted in the soil, three weeks before the first measurements at each site. Measurements were performed with an opaque survey chamber (Li-8100-104, LI-COR Biosciences, Nebraska, USA) connected to an LI-8100 analyzer (LI-COR Biosciences, Nebraska, USA). On each collar, the measurement period was set to 120 s the first 20 s of the measurement were considered dead-band, so the flux computation was limited to 80 s. See Montagnani et al. [8] for further details about the measurement settings. Starting on July 21, 2017, SR measurements were performed periodically, about once per month, until July 20, 2018, for a total of 17 measurement days. The first 4 measurement series were performed every three weeks at all the sites. During the winter period, SR measurements were performed only in the locations at a lower altitude because of snow in the high-elevation locations. The measurement calendar for the different sites is provided in S1. During measurements, the air temperature was measured inside the survey chamber (at 0.1 m above ground, RHT Plus, Skye Instruments, UK). A soil temperature profile was installed at control site B according to ICOS protocol [47]. Specifically, we used the -5 cm soil T data provided by a CS605 probe, Campbell Scientific, USA. We recorded soil temperature continuously at 30 min intervals for the whole experimental period (July 2017 - July 2018). In addition, during the period May-July 2018, we placed at all the sites iButton sensors (Maxim integrated, USA) at 5 cm below the soil surface.

### Soil sampling and analysis

At the end of the last measurement session (July 2018), leaf litter present in each collar was sampled. The soil in each collar was sampled using a 4.8 split-corer (Eijkelkamp, NL), until 20 cm depth. In the laboratory, soil samples were weighed and sieved at 2 mm mesh size to separate roots, stones, and coarse organic matter fragments. Collected leaf litter and fine roots (< 2 mm diameter) were weighted after oven-drying at 105 ± 5 °C. Soil bulk density was determined by dividing the weight of sieved soil by the core volume. Soil pH was measured using a pH-meter (CRISON pH-Meter Basic 20+Electrod: Hach 50 10T CRISON, Barcelona, Spain) in water. Root, litter, and soil samples were analyzed for organic C and N content using a FlashEA^™^ 1112 Elemental Analyzer (Thermo Fisher Scientific, Waltham, MA, USA).

### Data elaboration and statistical analysis

The mean amount (± SD) of accumulated C and N stock in the soil, root, and litter were computed for each site. Soil and C and N stocks were obtained as follows [48]:

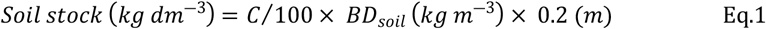

Where C is the mean soil organic C or total N content, *BD*_*soil*_ is the soil bulk density (kg dm^−3^) and 0.2 m is the sampling depth.

Root C and N stocks were determined with the same computational approach, using root density (kg dm^-3^) in place of soil BD and root C and N content in place of soil C and N content. Litter C and N amounts were obtained by multiplying the litter mass by C or N content and dividing by the collar area.

Soil respiration data collected from each measurement point (collar) were related to chamber air temperature using a logistic model [26]:

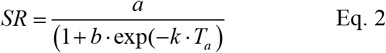

Where SR is soil respiration, a is the maximum value of SR, b determines the elongation of the SR curve along the x-axis, k is the logistic growth rate or steepness of the SR curve along the x-axis, Ta is air temperature. Furthermore, SR data were also fitted with a Q10 model [49,50]:

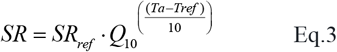

Where SR is the soil respiration, SR_ref_ is the fitted SR at the reference temperature of 10°C (T_ref_), Q10 is the temperature sensitivity of SR, defined as the factor by which soil respiration increases with a 10°C temperature increase, and Ta is chamber air temperature.

Models were fitted to SR data using the nls package in R software. Model fitness was evaluated based on Akaike’s Information Criterion (AIC), R-squared (R^2^), Mean Absolute Error (MAE), and Root Mean Squared Error (RMSE). The Q10 model was used to obtain the SR_ref_ and the Q10 value for every collar. Linear regression was used to compare the air temperature measured continuously in the reference plot (site B) and air temperature inside the chamber during SR measurements for each collar in each site. The obtained linear regression models were then used to predict chamber air temperature for the whole experimental period, for each collar, with a 30 min time resolution. Therefore, the predicted chamber air temperature was used to predict SR values simultaneously for the whole experimental period, based on the logistic models relating SR with chamber temperature. For some collars, it was not possible to obtain a good fitness of the SR data using the logistic model; for these collars, the prediction soil respiration data from temperature was performed using the Q10 model developed for the same collar. Finally, the total cumulative SR for the whole experimental period was determined for each collar at each site.

Soil respiration response to biological variables (soil C, root C, litter C, root dry weight, soil N, root N, litter N, litter dry weight) was examined using Spearman’s Correlation Test and linear mixed-effects models (LMMs) fitted by restricted maximum likelihood (REML). Before applying LMMs, to avoid statistical errors, Variance Inflation Factor (VIF) was determined for biological variables, and variables with high VIF values were excluded from the model assessment. LMMs were built using the lme4 R package [50,51,52]. The models consisted of both fixed and random effects: biological variables were considered as fixed effects, and sampling plots (collars) nested in each site were used in the random-effects formula. R^2^ was used to summarize model goodness-of-fit together with AIC [53,54]. Since computed R^2^ by LMMs are a pseudo-R^2^ and technically incorrect, the r2glmm R package was used to computing R^2^ [54].To exclude the confounding effect of temperature from LMMs and correlations tests, environmental variables were related to SR_ref_ instead of SR [20,26,55]. Furthermore, to assess GPP and SR’s correlation, tree height was used as a covariate in LMMs and Spearman test. Statistical comparisons of average soil C and N, root C and N, litter C and N, soil bulk density, soil pH, and soil respiration in the different sites were performed by the Kruskal-Wallis test (Dunn test, p < 0.05) for the non-normally distributed data, and a one-way ANOVA for the normally distributed data (Tukey test, p < 0.05). The normality of the data and homogeneity of variance was checked by the Shapiro–Wilk test and Levene’s test, respectively [56,57]. To check linearity changes of SR, Q10, SOM with elevation, linear and nonlinear polynomial regressions were applied between elevation and environmental variables (soil C and N, root C and N, litter C and N, soil bulk density, soil pH). The linearity changes of these variables with elevation were detected based on the lowest AIC and the highest R^2^. The association of Q10, soil C and soil N with environmental variables were determined using Spearman’s Correlation Test. All statistical analyses were performed using R version 3.6.0 ([59], www.r-project.org).

## Results

### Environmental factors variability along the altitudinal gradient

A significant difference in soil C stock was found only between site E (3891 ± 2756 g C m^-2^), at the lowest altitude, where the C stock was smaller in comparison to sites A, B, and C (Fig 2a). No significant differences were found for soil N stock in the different sites (Fig 2e). Root biomass and root C and N stocks in site A were significantly higher than other sites (Fig 2b, d, f) and litter mass in site B was significantly higher than site E (Fig 2h). However, the accumulated C and N in the litter were not significantly different along the altitudinal gradient (Fig 2c, g). Significant differences were found between pH values in the different sites: the lowest value of soil pH was measured in site B (3.8 ± 0.1) and the highest value in site E (5.9 ± 0.2, Fig 2i). The highest bulk density value was found in site D (1.01 ± 0.27 g cm^-3^, Fig 2i) and the lowest was found in site B (0.26 ± 0.16 g cm^-3^; Fig 2j).

**Fig 2.**
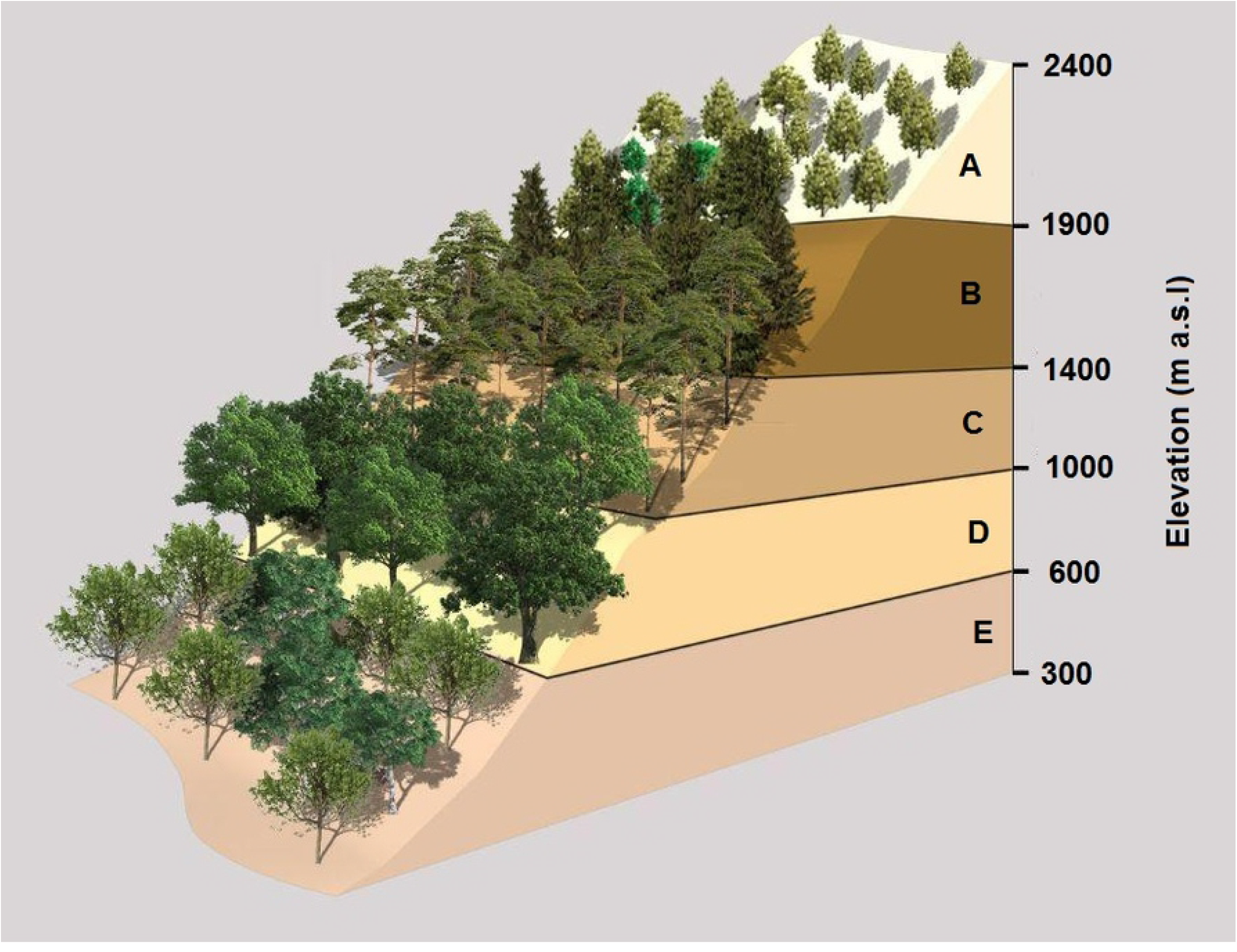
Stocks of (a), soil C (b,) root C (c), litter C (d), root dry weight (e,) soil N (f), root N (g), litter N (h), litter dry weight (i), soil pH, and soil bulk density (j), estimated in the different sites (A-E). Different lowercase letters indicate significant differences between sites according to the Kruskal-Wallis or ANOVA tests. Vertical bars represent the standard deviation of the mean for each site.

Based on the AIC and R^2^, the linear relationship with elevation appeared in model selection only for soil bulk density i.e., soil bulk density resulted to be linearly related to elevation (Table 2). On the contrary, soil C, soil N, fine root mass, and root C, root N, pH and litter mass data were fitted better with nonlinear relations with elevation (Table 2; more detail about equations can be found in Table S1; supplementary material). Furthermore, a significant negative correlation was found between soil C and soil N with soil pH, mean dominant tree height, and bulk density (Table 3).

**Table 2.**
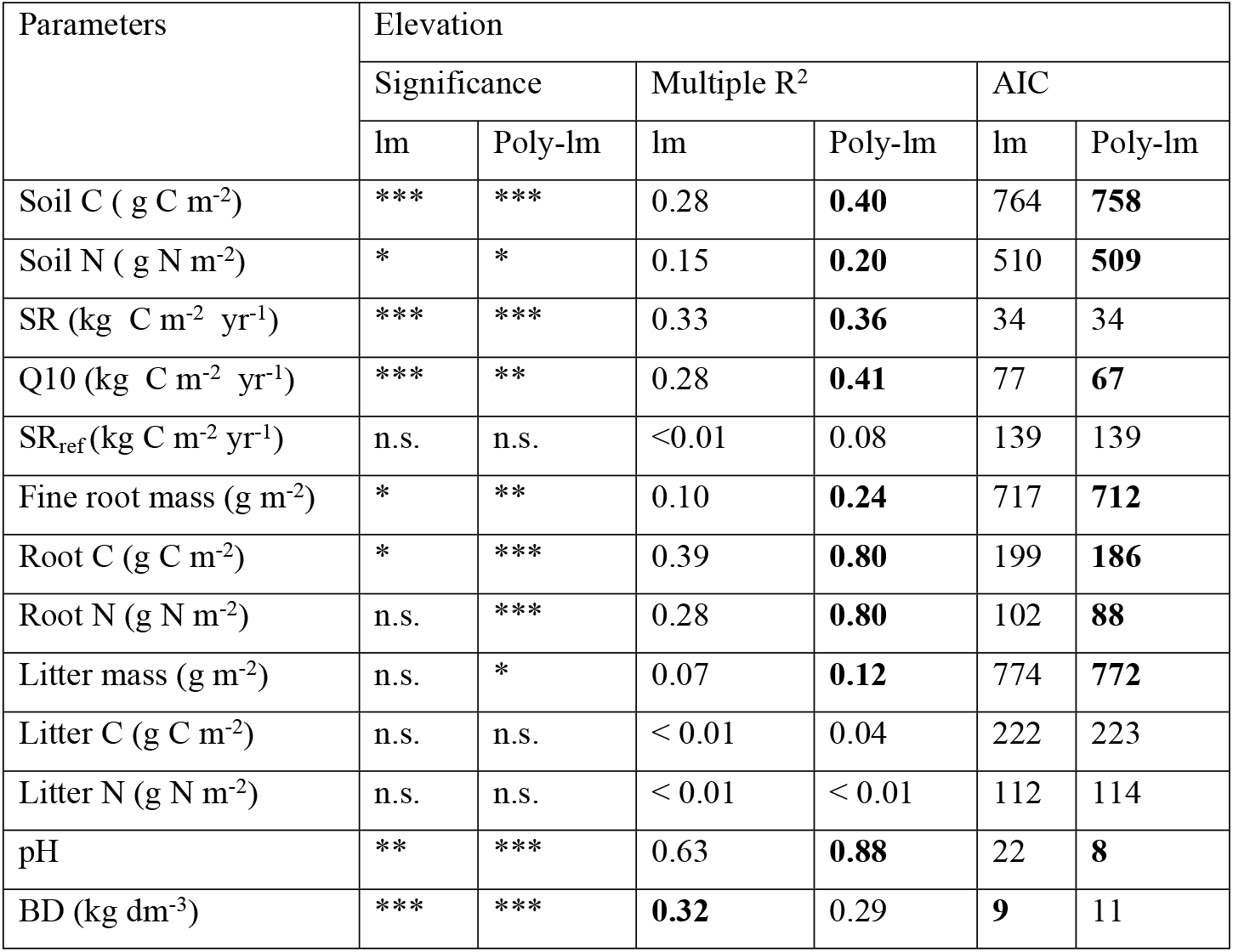
linear and polynomial regressions between the different parameters tested and elevation. Asterisks indicate significance levels: * - p ≤ 0.05, **- p ≤ 0.01, and *** - p ≤ 0.001, n.s. – nonsignificant. BD – soil bulk density; Q10 – temperature sensitivity of soil respiration; SR- cumulated soil respiration; SR_ref_ – respiration at the 10°C reference temperature; lm – linear regression model; Poly-lm – polynomial regression model. The model selection was based on the lowest AIC and the highest R^2^ (in bold).

**Table 3.**
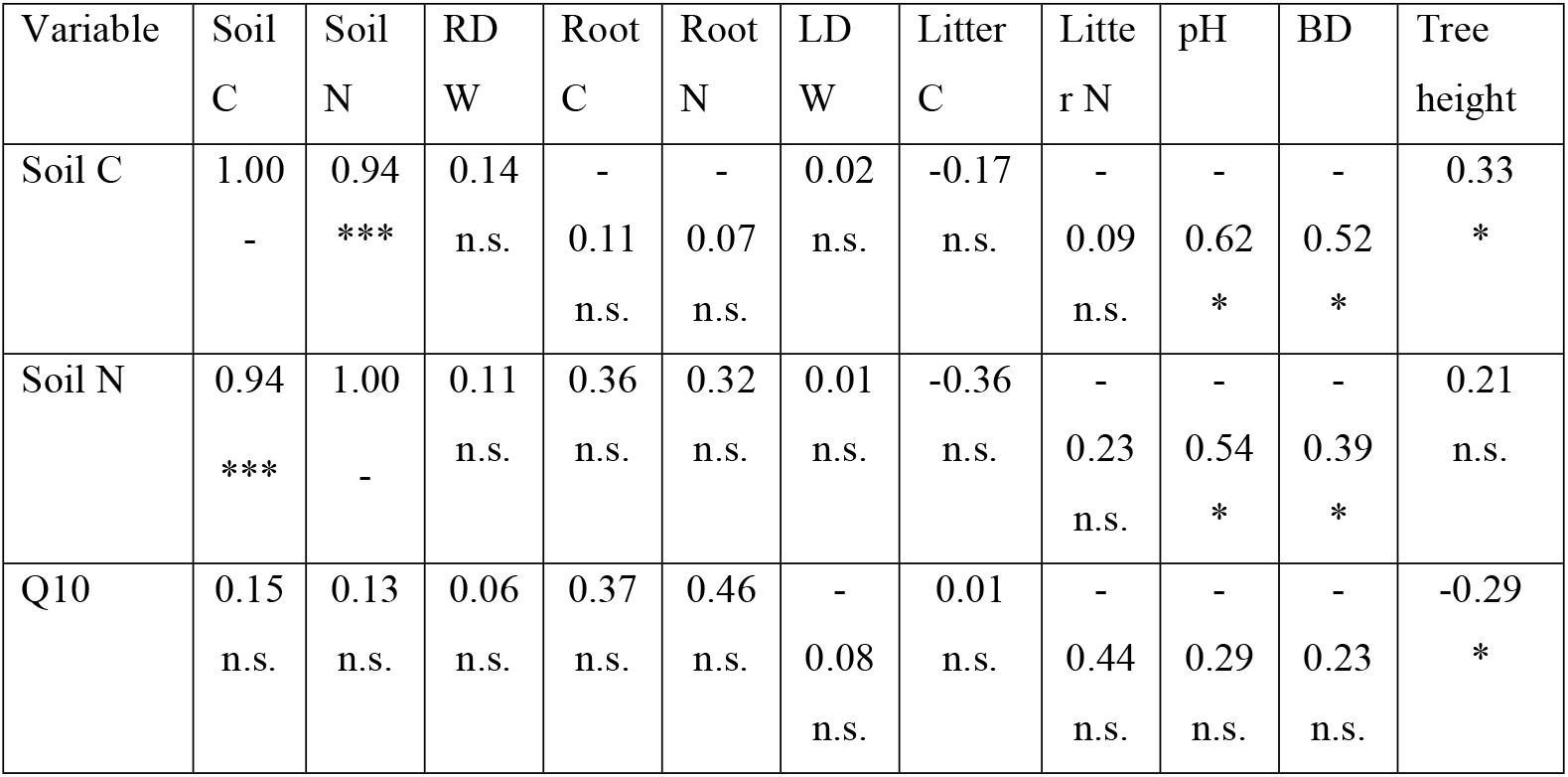
Spearman rank coefficients for the correlations between different variables (RDW – root dry weight, LDW – litter dry weight, BD – soil bulk density, Q10 – temperature sensitivity of soil respiration). Asterisks indicate significance levels: * - p ≤ 0.05, **- p ≤ 0.01, and *** - p ≤ 0.001; n.s. – nonsignificant.

### Soil respiration

Both logistic and Q10 models confirmed that soil respiration rates increased with temperature in all sites (Fig 3), and the seasonal pattern of SR was similar to that of air chamber temperature (S1 Fig).

**Fig 3.**
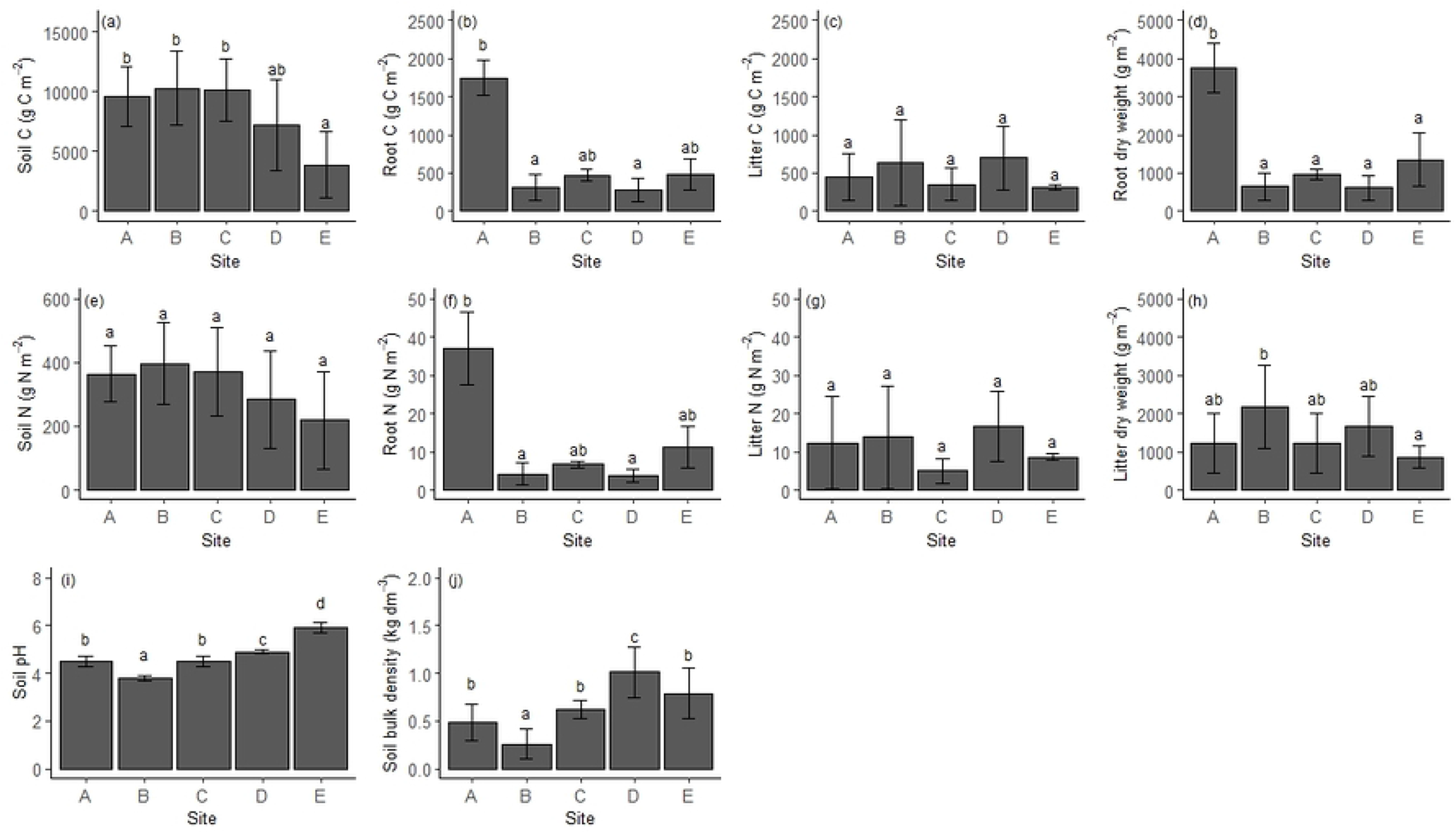
Rates of soil respiration against chamber air temperature in the different sites along the elevation gradient (A-E). The regression lines in the plots were built using the mean values of model parameters (R_ref_ and Q10 value for the Q10 model and a, b, and k value for the logistic model) obtained for different replicate collars of each site (n = 10).

A strong linear relationship was found between observed and predicted SR (R^2^ = 0.73; S2 Fig; supplementary material). Temperature explained between 55 % and 76% of the variance in soil respiration at the experimental sites (Table 4). The Q10 and SR_ref_ values obtained for the different sites ranged between 1.75 and 2.96, and between 2.17 and 4.49, respectively (Table 4).

**Table 4.** Mean values of Q10 (temperature sensitivity) and SR_ref_ (soil respiration at the temperature of 10°C) for each site; and the mean value of MAE (Mean Absolute Error), Root Mean Squared Error (RMSE), R^2^ (R-square), and AIC (Akaike information criterion) for each model (logistic and Q10). Different letters indicate significant differences between sites. Values are expressed as mean ± SD.

|  | A | B | C | D | E |
| --- | --- | --- | --- | --- | --- |
| Q10 | 2.96 ± 0.72<sup>b</sup> | 1.83 ± 0.25<sup>a</sup> | 1.90 ± 0.24<sup>a</sup> | 1.75 ± 0.15<sup>a</sup> | 1.75 ± 0.32<sup>a</sup> |
| $SR_{ref}$ | 2.17 ± 0.60<sup>a</sup> | 4.49 ± 0.71<sup>c</sup> | 2.24 ± 0.58<sup>a</sup> | 3.17 ± 1.03<sup>b</sup> | 2.98 ± 0.41<sup>ab</sup> |
| AIC logistic model | 32.95 ± 10.45 | 34.25 ± 6.45 | 34.19 ± 11.65 | 48.00 ± 14.55 | 49.76 ± 6.70 |
| AIC Q10 model | 30.42 ± 13.69 | 34.07 ± 7.31 | 35.39 ± 11.53 | 48.44 ± 14.13 | 49.11 ± 6.60 |
| $R^2$ logistic model | 0.76 ± 0.21 | 0.75 ± 0.20 | 0.75 ± 0.16 | 0.67 ± 0.11 | 0.58 ± 0.10 |
| $R^2$ Q10 model | 0.75 ± .23 | 0.71 ± 0.23 | 0.68 ± 0.21 | 0.61 ± 0.11 | 0.55 ± 0.11 |
| MAE logistic model | 1.06 ± 0.65 | 1.06 ± 0.45 | 0.68 ± 0.32 | 1.26 ± 0.62 | 1.29 ± 0.25 |
| MAE Q10 model | 1.07 ± 0.68 | 1.14 ± 0.52 | 0.79 ± 0.36 | 1.46 ± 0.76 | 1.39 ± 0.26 |
| RMSE logistic model | 1.36 ± 0.94 | 1.27 ± 0.50 | 0.86 ± 0.44 | 1.70 ± 0.89 | 1.71 ± 0.36 |
| RMSE Q10 model | 1.38 ± 1.0 | 1.36 ± 0.55 | 0.99 ± 0.49 | 1.87 ± 1.02 | 1.77 ± 0.38 |

The Q10 value recorded in site A (highest elevation) was significantly different from the others (Table 4). A significant linear relationship was identified between Q10 and elevation (Table 2). However, the trend of Q10 against temperature was better described by a nonlinear relation (Table 2). Furthermore, a significant negative correlation was found between Q10 and mean dominant tree height (Table 3). No significant relationship between SR_ref_ and altitude was found (Table 2). However, significant differences were found between experimental sites, as the highest SR_ref_ value was recorded in site B and the lowest values in site A, C, and E (Table 4).

The cumulative SR in site A was significantly lower than in the other sites (Fig 4). The results of the statistical analysis also confirmed a nonlinear relationship between cumulative SR and elevation (Table 2).

**Fig 4.**
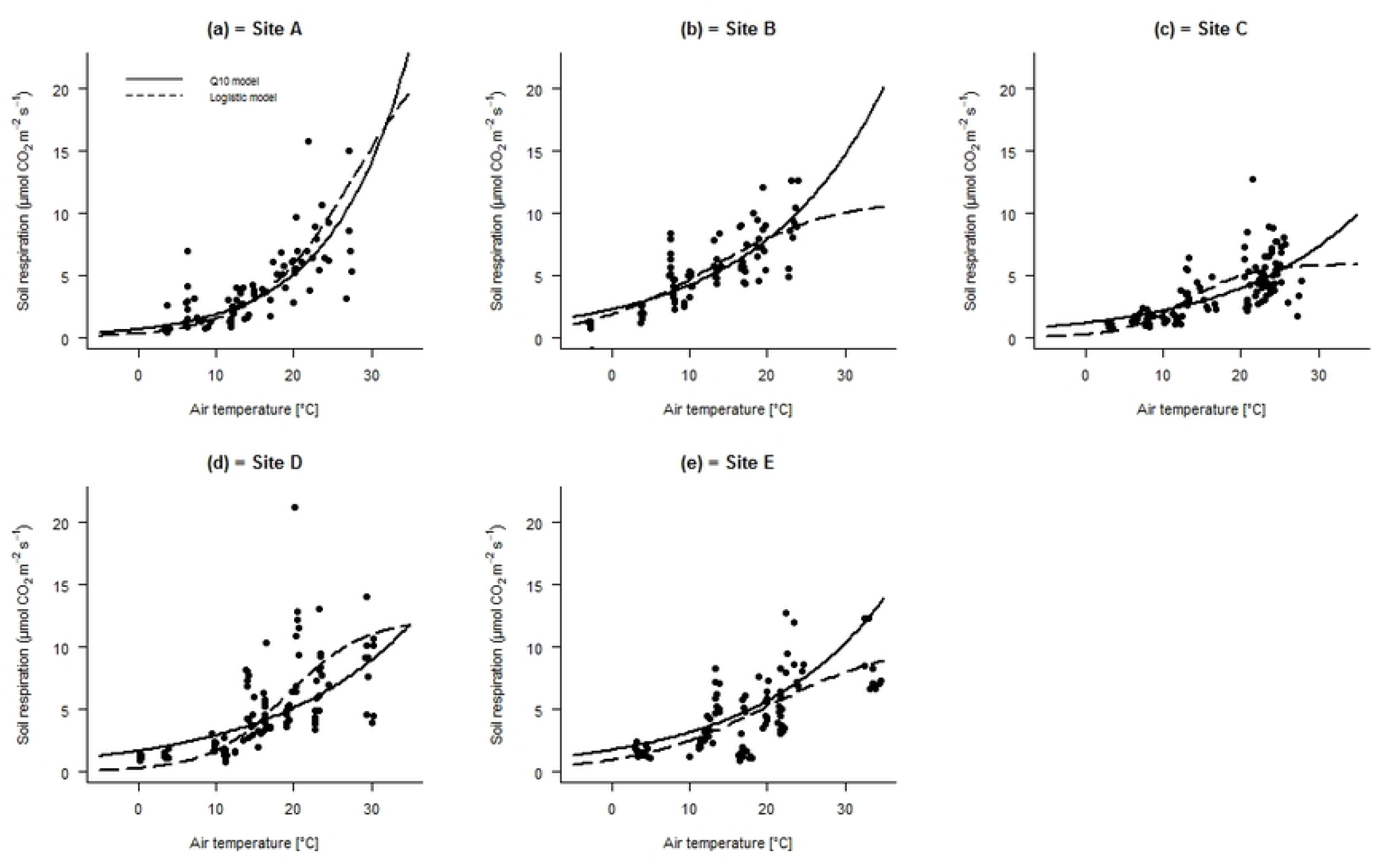
Total cumulative soil respiration (kg C m^-2^ yr^-1^) calculated for the different sites (A-E). Values are indicated on the bars. Error bars indicate standard deviation. Different letters on the bars indicate significant differences between sites according to the ANOVA test.

Soil C, mean dominant tree height, and litter dry weight resulted in the best variables to explain SR (SR_ref_; at 10 C) in LMMs (VIF<10; Table 5). According to the model, about 0.22 of SR was explained by tree height (R^2^ = 0.22, Table 5). Meanwhile, a positive association between SR and mean dominant tree height and a negative association between SR and root C and N were found by Spearman’s Correlation Test (Fig 5).

**Table 5.** Results of linear mixed-effects models testing biological variables impact on the SR_ref_; at 10 C. VIF – Variance Inflation Factor. Parameters in bold show significant correlations.

| Model variables | Value | VIF | p-value | $R^2$ |
| --- | --- | --- | --- | --- |
| Intercept | 1.61 |  | 0.003 | 0.34 |
| Soil C | < - 0.01 | 1.20 | 0.92 | < 0.01 |
| **Tree height** | **0.06** | **1.28** | **0.002** | **0.22** |
| Litter dry weight | < 0.01 | 1.07 | 0.14 | 0.06 |

**Fig 5.**
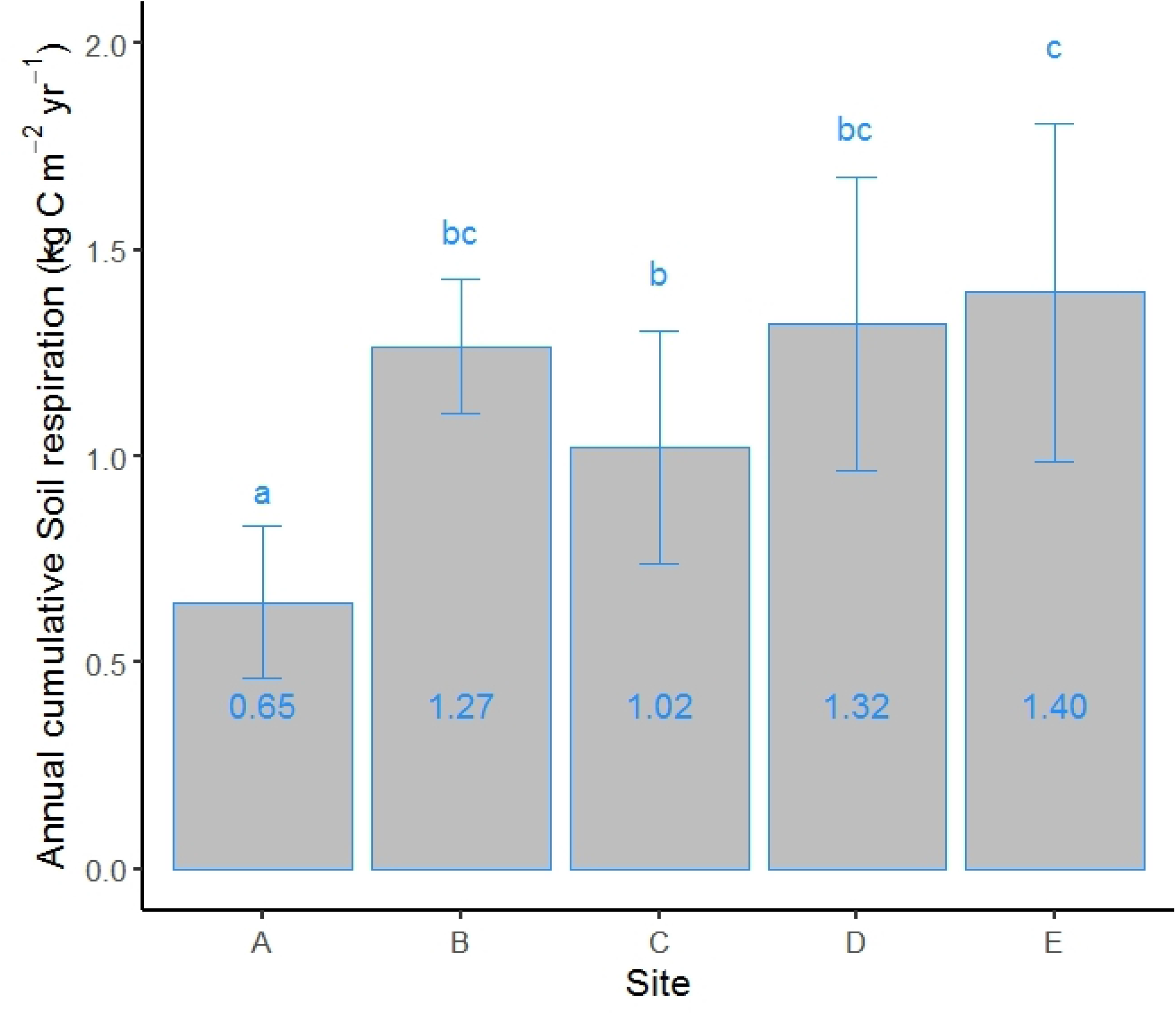
Relations of reference soil respiration at 10°C (SR_ref_) and different site properties.

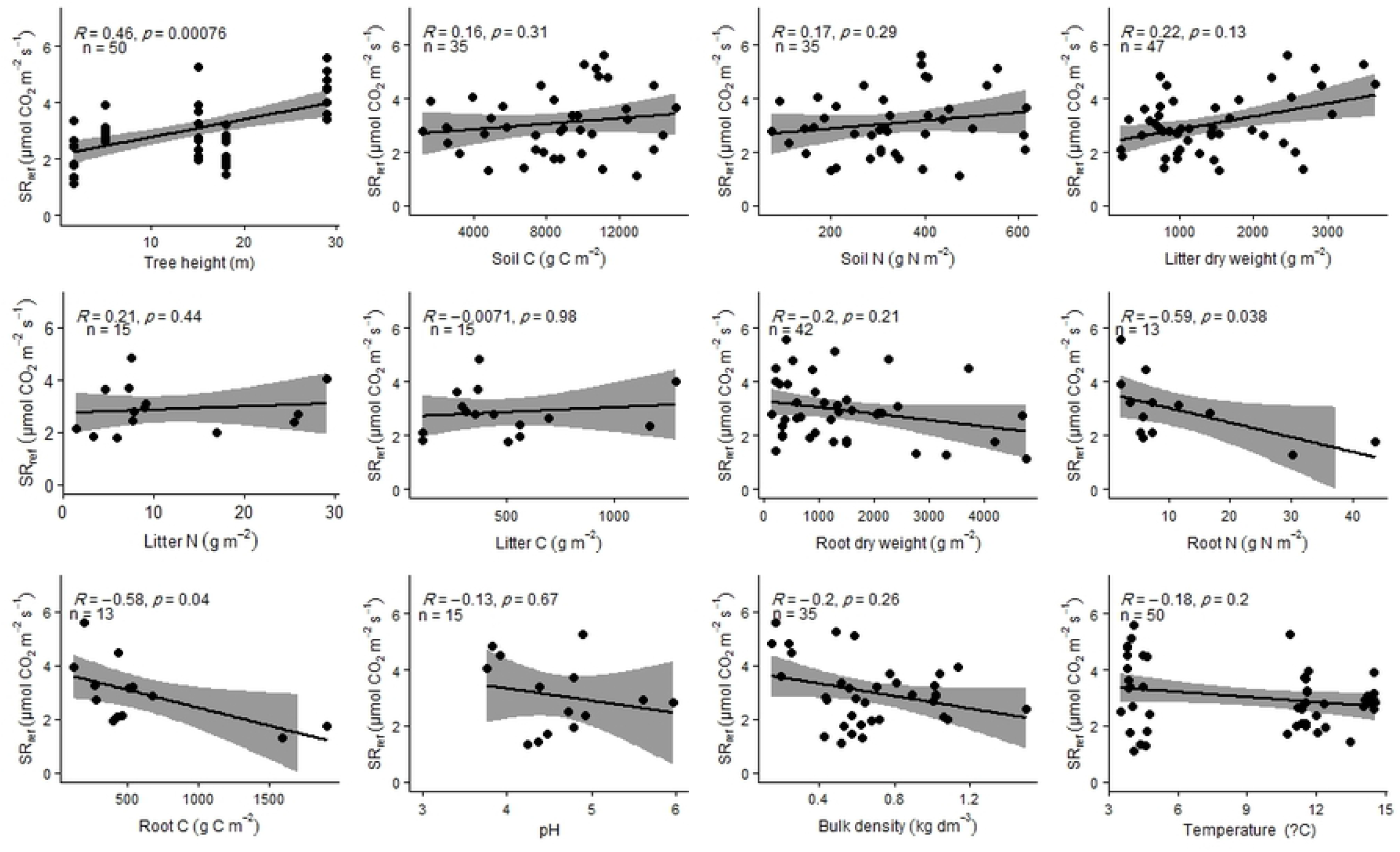

## Discussion

### Organic C and N content along the altitudinal gradient

Overall, our data confirms an increase of the soil organic C stock in soil with increasing elevation [7,24, 25,27,59,60,61,62]. The low temperature can limit the decomposition of organic matter at high altitudes, as the temperature is the main driver for the loss of the soil organic C. For this reason, altitude could induce a significant increase in SOM [23,59,62,63,63]. However, SOM increment was not linear along the elevation gradient and therefore, our first hypothesis was not confirmed. The Nonlinear relationship of SOM with elevation and the high value of soil C in site C (with a mean temperature of 12 C) suggests that other factors other than temperature have influenced SOM accumulation. Different microclimatic or micromorphological conditions caused by differences in slope and aspect can influence C storage in soils [33,35,65]. However, in the present study, all the sites are characterized by similar slopes and by the same south or south-east facing; therefore, we tend to exclude an influence of these factors on soil C accumulation in the examined sites. According to recent studies, SOM is not consistently related to variation in climatic conditions along elevation gradients; it is also strongly affected by productivity or by vegetation type/composition [30,33,66,67]. Our data confirms a significant positive correlation between mean dominant tree height and soil C (Table 3). Therefore, the higher amount of soil C found in the present study at intermediate altitude could be explained by higher site productivity in sites B and C in particular, which is also suggested by the mean dominant tree height.

Soil pH and bulk density are considered two of the main variables influencing other soil properties, soil microbial activity and soil respiration [68,69]. Generally, at high elevation, the higher precipitation and lower evapotranspiration rate decrease soil pH by increasing the leaching of basic cations [65,70,71,72,73]. This is confirmed by the strong relationship between elevation and soil pH found in the present study (Table 2, Fig 2). In addition, soil bulk density was significantly diminished by increasing elevation (Fig 2; Table 2). One of the main factors affecting soil bulk density is SOM content [74]. Therefore, the lowest values of soil bulk density at high elevation could be explained by the high amount of soil C, as confirmed by the negative association found between soil C and soil bulk density (Table 3), previously reported in other studies [7, 61,62,75].

### Factors affecting soil respiration

The total SR observed in the sites is within the range reported for similar forests [26,60]. The decrease in SR along the elevation gradient observed in the present study could be explained by the reduction of temperature along elevation. In fact, temperature resulted in the main controlling factor on SR, explaining most of the variability of SR. This result is in agreement with other studies performed along altitudinal gradients, reporting that temperature can explain between 55% and 76% of the SR variability [6, 9, 14,27,63, 76]. On the other hand, the annual cumulative SR in site B was 2 times larger than in site A, which has the same mean annual temperature. This finding, together with nonlinear SR concentrations with elevation/temperature, suggests that other environmental factors can have a role in regulating SR [34,60]. For instance, Grand et al. [36] reported that soil and vegetation heterogeneity strongly affect soil carbon efflux in complex geomorphic terrain. In the present study, all five sites were established on relative homogeneity of soil substratum; therefore a high SR rate in site B could not be resulted by the confounding role of the soil parent material. Site B is an uneven-aged dense forest stand, and the mean dominant tree height is approximately 29 m (Table 1). Since tree height can be used as a proxy of gross primary production (GPP), the high SR rate in site B could be attributed to high GPP [9,29,66,77,78,79], which can provide substrates for root and microbial respiration through photosynthesis [66,80]. This finding is confirmed by LMMs and correlation test evidencing a significant positive relation between SR_ref_ and mean dominated tree height, therefore indicating that, after removing the effect of temperature, productivity results in one of the main factors affecting SR.

At a global scale, SR has been related to soil C, litter production and pH, and negatively correlated with soil bulk density; therefore, a high value of soil C and litter accumulation could lead to an increase in soil respiration [20,66,81,82,83]. In the present study, the highest amount of soil C and dry litter weight were also observed in site B. However, we could not find a significant correlation between SR and soil C and litter dry weight (Fig 5; Table 5).

An increase in SR has also been observed as a consequence of increasing soil pH between 4 and 7, because of the positive effect of pH on soil microbial activity within this range [1,68,84,85,86]. In contrast, SR and bulk density are generally negatively correlated, as a low SR indicates increasing rates of SOM accumulation and therefore a decrease in bulk density [82]. Furthermore, SR declines with increasing bulk density due to the lower soil porosity and oxygen availability for microbial activity in compacted soils [18,59,87]. However, our analysis did not confirm a significant correlation of SR with pH and bulk density (Table 5; Fig 5). Prediction of SR is difficult because of a range of factors such as aspect, slope, and soil types [35,36,60,65]. In the present study, by minimizing the confounding role of these parameters, we conclude that the most important controlling factors on SR along an Alpine elevation gradient were temperature and vegetation type/composition or GPP.

### Temperature sensitivity of soil respiration (Q10)

The temperature sensitivity of SR is an important ecological model parameter, and according to previous studies, its value is mainly controlled by temperature [14,17,88,89]. The Q10 and SRref values found at site B substantially confirm the values found by Acosta et al. [90] at the same site (2.0 and 4.09, respectively). Although at a smaller spatial scale, Acosta et al. [90] also found an increasing SRref as a function of stand age (and consequently height). The significant trend of Q10 with elevation in this research confirms the results of previous studies and the higher sensitivity of high elevation ecosystems to global warming [14,91,92,93,94]. However, the only significant difference was found between the Q10 value at site A (higher elevation) and the other sites (Table 4). The Q10 value in site A was also significantly higher than site B, characterized by a similar mean temperature. According to different studies, Q10 is negatively related to pH and positively dependent on soil C [55,95]. However, the amount of soil C and pH could not be the reasons for the lower value of Q10 in site B, which is characterized by a lower pH value and by a similar amount of soil C. Temperature sensitivity of SR can also be affected by forest structure [96]. Dense forest stands with a closed canopy can create a specific understory microclimate by providing a cool shelter during heat waves, which can decrease daily maximum air temperature by up to 5.1 °C [97,98]. Therefore, we can hypothesize that the dense forest stand site B, with the highest mean dominant height, is less sensitive to global warming. This feature was also described by Niu et al. [99] for the same site and confirmed by a significant negative correlation between Q10 and mean dominating tree height (Table 3). Whether a sylviculture that maintains high and old forests with a complex structure also achieves a lower sensitivity to climate change by reducing the depletion of C stored in the soil, is a novel research question emerging from this research. If the findings obtained here will be confirmed, it would imply that more conservative forest management can not only maintain current C stocks in the biomass but also leads to a reduced sensitivity to the temperature of the C stored in the soil.

## Conclusions

In this study, a significant nonlinear relationship between SR, SOM, Q10, and elevation was detected along the examined Alpine altitudinal gradient, rejecting our initial linearity hypothesis. Our data confirmed a negative trend between SR and elevation. On the contrary, SOM and Q10 showed a positive trend with elevation. These results lead us to conclude that temperature is the major controlling factor on the annual soil respiration, Q10, and SOM, but its regulating role may be strongly affected by site biological characteristics, particularly by GPP or vegetation type/composition. The high value of Q10 detected at high elevation confirmed a higher potential vulnerability of high mountain ecosystems to climate change, where small temperat
ure changes can induce stronger increase CO2 emissions. However, the site with the highest tree age and height and more complex structure showed high SRref and moderate Q10, indicating that the length in the life cycle and related changes in forest structure can dampen, to some extent, the effects of climate change on ecosystems and decrease the positive feedback due to soil CO2 emissions to the atmosphere.

## Supporting Information

**S1**. **Collected soil respiraton data by soil chambers**. Used for soil respiration modeling

**S2**. **Measured soil physicochemical properties**. Used for the analysis.

**S1 Fig**. **Predicted daily mean air temperature (°C**, **closed circles) and soil CO**_**2**_ **fluxes (µmol CO**_**2**_ **C m**^**-2**^ **s^-1^**, **open circles) during the experimental period (July 2017- July 2018)**. Mean annual air temperature (Tair, °C) and total cumulative soil respiration in the entire experimental period (SR, kg C m^-2^) are indicated for each site on the top of the relative plots.

**S2 Fig**. **Observed soil respiration vs**. **predicted soil respiration by model**.

**S1 Table**. **The developed equations by linear and polynomial regressions between the different parameters tested and elevation**. BD – soil bulk density; Q10 – temperature sensitivity of soil respiration; SR- cumulated soil respiration; SRref – respiration at the 10°C reference temperature; Elev – elevation.

## References

1. Luo Y, Zhou X. Soil respiration and the environment. Sandiego, CA: Academic Press; 2006.

2. Valentini R, Matteucci G, Dolman AJ, Schulze ED, Rebmann C, Moors EJ, et al. Respiration as the main determinant of carbon balance in European forests. Nature. 2000; 404: 861–865. DOI: 10.1038/3500908.

3. Scharlemann JP, Tanner EV, Hiederer R, Kapos V. Global soil carbon: understanding and managing the largest terrestrial carbon pool. Carbon Manag. 2014; 5: 81–91.

4. Makita N, Kosugi Y, Sakabe A, Kanazawa A, Ohkubo S, Tani M. Seasonal and diurnal patterns of soil respiration in an evergreen coniferous forest: Evidence from six years of observation with automatic chambers. PLoS ONE, 2018; 13(2): e0192622. https://doi.org/10.1371/journal. pone.0192622

5. Cong WF, Van Ruijven J, Mommer L, De Deyn GB, Berendse F, Hoffland E. Plant species richness promotes soil carbon and nitrogen stocks in grasslands without legumes. Journal of Ecology. 2014; 102(5): 1163–1170. DOI 10.1111/1365-2745.12280.

6. Luo W, Li MH, Sardans J, XT Lü, Wang C, Peñuelas J, Wang Z, Han XG, Jiang Y. Carbon and nitrogen allocation shifts in plants and soils along aridity and fertility gradients in grasslands of China. Ecol Evol. 2017; 7(17): 6927–6934, DOI 10.1002/ece3.3245.

7. Devi SB, SSSS Sherpa. Soil carbon and nitrogen stocks along the altitudinal gradient of the Darjeeling Himalayas, India. Environ Monit Assess. 2019; 191 (361): https://doi.org/10.1007/s10661-019-7470-8

8. Montagnani L, Badraghi A, Speak AF, Wellstein C, Borruso L, Zerbe S, Zanotelli D. Evidence for a non-linear carbon accumulation pattern along an Alpine glacier retreat chronosequence in Northern Italy. PeerJ. 2019; 7:e7703, DOI 10.7717/peerj.7703.

9. Janssens I.A, Lankreijer H, Matteucci G, Kowalski AS, Buchmann N, Epron D. Productivity overshadows temperature in determining soil and ecosystem respiration across European forests. Glob Chang Biol. 2001; 7: 269–278.

10. Hopkins F, Gonzalez-Meler MA, Flower CE, Lynch DJ, Czimczik C, Tang J, Subke JA. Ecosystem-level controls on root-rhizosphere respiration, New Phytol. 2013; 199: 339e351, http://dx.doi.org/10.1111/nph.12271.

11. Scandellari F, Zanotelli D, Ceccon C, Bolognesi M, Montagnani L, Cassol P, Melo GW, Tagliavini M. Enhancing prediction accuracy of soil respiration in an apple orchard by integrating photosynthetic activity into a temperature-related model, Eur J Soil Biol. 2015; 70: 77–87. doi: 10.1016/j.ejsobi.2015.07.006.

12. Zhang ZS, Dong XJ, Xu BX, Chen YL, Zhao Y, Gao YH, Hu YG, Huang L. Soil respiration sensitivities to water and temperature in a revegetated desert. J. Geophys. Res. Biogeosci. 2015; 120: 773–787. doi:10.1002/2014JG002805

13. Chen D, Yu M, González G, Zou X, Gao Q. Climate Impacts on Soil Carbon Processes along an Elevation Gradient in the Tropical Luquillo Experimental Forest. Forests. 2017. 8; 90: doi:10.3390/f8030090

14. Ma M, Zang Z, Xie Z, Chen Q, Xu W, Zhao C, Shen G. Soil respiration of four forests along elevation gradient in northern subtropical China. Ecol Evol. 2019; 9: 12846–12857.

15. Reichstein M, Beer C. Soil respiration across scales: the importance of a model-data integration framework for data interpretation. J Plant Nutr Soil Sci. 2008; 171: 344e354.

16. Subke JA, Bahn M. On the ‘temperature sensitivity’ of soil respiration: Can we use the immeasurable to predict the unknown? Soil Biol Biochem. 2008; 42 (9): 1653–1656.

17. Lloyd J, Taylor JA. On the temperature dependence of soil respiration, Funct Ecol. 1994; 8(3): 315–323. doi:10.2307/2389824.

18. Xu M, Qi Y. Soil surface CO2 efflux and its spatial and temporal variation in a young ponderosa pine plantation in California. Global Change Biol. 2001; 7: 667 – 677. https://doi.org/10.1046/j.1354-1013.2001.00435.x

19. Janssens IA, Dore S, Epron D, Lankreijer H, Buchmann N, Longdoz B, Brossaud J, Montagnani L. Climatic influences on seasonal and spatial differences in soil CO2 efflux. In: Valentini R (ed) Fluxes of carbon, water and energy of European forests. Berlin: Springer; 2003.

20. Reichstein M, Rey A, Freibauer A, Tenhunen J, Valentini R, Banza R. Modeling temporal and large-scale spatial variability of soil respiration from soil water availability, temperature and vegetation productivity indices. Global Biogeochem Cycles. 2003; 17: doi: 10.1029/2003GB002035. Van’t Hoff JH. Lectures on theoretical and physical chemistry. In Chemical Dynamics Part I (pp. 224–229). London: Edward Arnold.

21. Vereecken, H., Pachepsky, Y., Simmer, C., Rihani, J., Kunoth, A., Korres, W., et al., 2016. On the role of patterns in understanding the functioning of soil-vegetation atmosphere systems. J Hydrol. 1898; 542: 63–86. https://doi.org/10.1016/j.jhydrol.2016.08.053

22. Lomolino MV. Elevation gradients of species-density: historical and prospective views. Glob Ecol Biogeogr. 2001; 10: 3–13.

23. Prietzel J, Zimmermann L, Schubert A, Christophel D. Organic matter losses in German Alps forest soils since the 1970s most likely caused by warming. Nat Geosci. 2016; 1–8, DOI: 10.1038/NGEO2732

24. Shedayi AA, Xu M, Naseer L, Khan B. Altitudinal gradients of soil and vegetation carbon and nitrogen in a high altitude nature reserve of Karakoram ranges. SpringerPlus. 2016; 5: 320. DOI 10.1186/s40064-016-1935-9

25. Jiang L, He Z, Liu J, Xing C, Gu X, Wei C, Zhu J, Wang X. Elevation Gradient Altered Soil C, N, and P Stoichiometry of Pinus taiwanensis Forest on Daiyun Mountain. Forests. 2019; 10: 1089. doi:10.3390/f10121089

26. Rodeghiero M, Cescatti A. Main determinants of forest soil respiration along an elevation/temperature gradient in the Italian Alps. Glob Chang Biol. 2005; 11: 1024–1041. doi: 10.1111/j.1365-2486.2005.00963.x

27. Shi Z, Wang JS, He R, Fang YH, Xu ZK, Quan W, Zhang ZX, Ruan HH. Soil respiration and its regulating factor along an elevation gradient in Wuyi Mountain of Southeast China. Chinese J Ecol. 2008; 27 (4): 563–568.

28. Luo S, Liu G, Li Z, Hu C, Gong L, Wang M, Hu H. Soil respiration along an altitudinal gradient in a subalpine secondary forest in China. iForest. 2014; 8: 526–532.

29. Kane ES, Valentine DW, Schuur EAG, Dutta K. Soil carbon stabilization along climate and stand productivity gradients in black spruce forests of interior Alaska. Can. J. For. Res. 2005; 35: 2118–2129.

30. Djukic L, Zehetner F, Tatzber M, Gerzabek MH. Soil organic-matter stocks and characteristics along an Alpine elevation gradient. J Plant Nutr Soil Sci. 2010; 173: 30–38. DOI: 10.1002/jpln.200900027

31. Kunkel ML, Flores AN, Smith TJ, McNamara JP, Benner SG. A simplified approach for estimating soil carbon and nitrogen stocks in semi-arid complex terrain. Geoderma. 2011; 165 (1): 1–11.

32. Tian Q, He H, Cheng W, Bai Z, Wang Y, Zhan X. Factors controlling soil organic carbon stability along a temperate forest altitudinal gradient. Sci Rep. 2016; 6: 18783. DOI: 10.1038/srep18783

33. Garcia-Pausas J, Casals P, Camarero L, Huguet C, Sebastia MT, Thompson R, Romanya J. Soil organic carbon storage in mountain grasslands of the Pyre-nees, effects of climate and opography. Biogeochemistry. 2007; 82: 279–289. https://doi.org/10.1007/s10533-007-9071-9.

34. Zimmermann M, Meir P, Bird MI, Malhi Y, Ccahuana AJQ. Temporal variation and climate dependence of soil respiration and its components along a 3000 m altitudinal tropical forest gradient. Global Biogeochem Cy. 2010; 24: GB4012. doi: 10.1029/2010GB003787

35. Kobler J, Zehetgruber B, Jandl R, Dirnböck T, Schindlbacher A. Effects of slope aspect and site elevation on seasonal soil carbon dynamics in a forest catchment in the Austrian Limestone Alps. 19th EGU General Assembly, EGU2017, proceedings from the conference held. 2017; 23-28 April, in Vienna, Austria, p.16691

36. Grand S, Rubin A, Verrecchia EP, Vittoz P. Variation in Soil Respiration across Soil and Vegetation Types in an Alpine Valley. PLoS ONE. 2016; 11 (9): e0163968. https://doi.org/10.1371/journal.pone.0163968

37. Migliavacca M, Reichstein M, Richardson AD, Colombo R, Sutton MA, Lasslop G. Semiempirical modeling of abiotic and biotic factors controlling ecosystem respiration across eddy covariance sites. Global Change Biology. 2011; 17(1): 390–409. DOI: 10.1111/j.1365-2486.2010.02243.x.

38. Bréchet L, Ponton S, Alméras T, Bonal D, Epron D. Does spatial distribution of tree size account for spatial variation in soil respiration in a tropical forest?. Plant Soil. 2011; 347 (293): https://doi.org/10.1007/s11104-011-0848-1

39. Xu X, Shi Z, Li D, Zhou X, Sherry RA, Luo Y. Plant community structure regulates responses of prairie soil respiration to decadal experimental warming. Glob Change Biol. 2015; 21: 3846–3853. https://doi.org/10.1111/gcb.12940

40. Tian Q, Wang D, Tang Y, Li Y, Wang M, Liao C, Liu F. Topographic controls on the variability of soil respiration in a humid subtropical forest. Biogeochemistry. 2019; 145: 177– 192. https://doi.org/10.1007/s10533-019-00598-x

41. Tonon G, Dezi S, Ventura M, Scandellari F. The Effect of Forest Management on Soil Organic Carbon. In: Sauer TJ, Eiler JM, Sivakumar MVK (eds) Sustaining Soil Productivity in Response to Global Climate Change: Science, Policy, and Ethics. John Wiley & Sons, Inc. 2011; pp 225–238

42. Greiser C, Meineri E, Luoto M, Ehrlén J, Hylander K. Monthly microclimate models in a managed boreal forest landscape. Agri For Meteorol. 2018; 250–251: 147–158. https://doi.org/10.1016/j.agrformet.2017.12.252.

43. Carey JC, Tang J, Templer PH, Kroeger KD, Crowther T, et al. Temperature response of soil respiration largely unaltered with experimental warming. PNAS. 2016; 113 (48): 13797– 13802.

44. Tang J, Cheng H, Fang C. The temperature sensitivity of soil organic carbon decomposition is not related to labile and recalcitrant carbon. PLoS ONE. 2017; 12 (11): e0186675. https://doi.org/10.1371/journal.pone.0186675

45. Montagnani L, Manca G, Canepa E, Georgieva E, Acosta M, Feigenwinter C, Janous D. A new mass conservation approach to the study of CO2 advection in an alpine forest. J. Geophys. Res. Atmos. 2009; 114: D07306, DOI:10.1029/2008JD010650.

46. Xu X, Yi C, Montagnani L, Kutter E. Numerical Study of the Interplay between Thermo-topographic Slope Flow and Synoptic Flow on Canopy Transport Processes, Agric For Meteorol. 2018; 255: 3–16. https://doi.org/10.1016/j.agrformet.2017.03.004.

47. Gielen B, Acosta M, Altimir N, Buchmann N, Cescatti A, Ceschia E, Fleck S. Soil-meteorological measurements at ICOS monitoring stations in terrestrial ecosystems. Int Agrophys. 2019; 32: 645–664. DOI: 10.1515/intag-2017-0048

48. Ventura M, Panzacchi P, Muzzi E, Magnani F, Tonon G. Carbon balance and soil carbon input in a poplar short rotation coppice plantation as affected by nitrogen and wood ash application. New Forests. 2019. https://doi.org/10.1007/s11056-019-09709-w

49. Richards FJ. A flexible growth function for empirical use. J Exp Bot. 1959; 10: 290 – 300.

50. Janssens IA, Pilegaard K. Large seasonal changes in Q10 of soil respiration in a beech forest. Glob Chang Biol. 2003; 9: 911–918.

50. Crawley M.J. The R Book. Chichester: John Wiley & Sons, Ltd.; 2007.

51. Zuur A, Ieno EN, Walker N, Saveliev AA, Smith GM. Mixed Effects Models and Extensions in Ecology with R. New York: Springer; 2009.

52. Bates D, Maechler M, Bolker B, Walker S. lme4: linear mixed-effects models using Eigen and S4. R package version 1.1-7; 2014.

53. Nakagawa S, Holger S. A general and simple method for obtaining R2 from generalized linear mixed effects models. Methods Ecol Evol. 2013; 4: 133–142.

54. Jaeger BC, Edwards LJ, Das K, Sen PK. An R2 Squared Statistic for Fixed Effects in the Generalized Linear Mixed Model. J Appl Stat. 2016; 44 (6): 1086–1105

55. Meyer N, Welp G, Amelung W. The temperature sensitivity (Q10) of soil respiration: Controlling factors and spatial prediction at regional scale based on environmental soil classes. Global Biogeochem Cy. 2018; 32: 306–323. https://doi.org/10.1002/2017GB005644

56. Shapiro SS, Wilk MB. Analysis of variance test for normality. Biometrika. 1965; 52: 591–611.

57. Levene H. In Contributions to Probability and Statistics: Essays in Honor of Harold Hotelling, I. Olkin et al. eds., Stanford University Press, 1960; 278–292.

58. R Core Team. R: A language and environment for statistical computing. R Foundation for Statistical Computing, Vienna, Austria. 2019. Available online at https://www.R-project.org/.

59. Wang G, Zhou Y, Xu X, Ruan H, Wang J. Temperature Sensitivity of Soil Organic Carbon Mineralization along an Elevation Gradient in the Wuyi Mountains, China. PLoS ONE. 2013; 8(1): e53914. doi:10.1371/journal.pone.0053914

60. Chatterjee A, Jenerette GD. Variation in soil organic matter accumulation and metabolic activity along an elevation gradient in the Santa Rosa Mountains of Southern California, USA. J Arid Land. 2015; 7(6): 814–819. doi: 10.1007/s40333-015-0085-1

61. Tsozuéa D, Nghonda JP, Tematio P, Basga SD. Changes in soil properties and soil organic carbon stocks along an elevation gradient at Mount Bambouto, Central Africa. Catena. 2019; 175: 251–262. https://doi.org/10.1016/j.catena.2018.12.028

62. de la Cruz-Amo L, Bañares-de-Dios G, Cala V, Granzow-de la Cerda I, Espinosa CI, Ledo A, Salinas N, Macía M.J, Cayuela L. Trade-offs among aboveground, belowground, and soil organic carbon stocks along altitudinal gradients in Andean Tropical Montane Forests. Front. Plant Sci. 2020; 11:106. doi: 10.3389/fpls.2020.00106

63. Kirschbaum MUF. The temperature dependence of organic-matter decomposition-still a topic of debate. Soil Biol Biochem. 2006; 38: 2510–2518

64. He X, Hou E, Liu Y, Wen D. Altitudinal patterns and controls of plant and soil nutrient concentrations and stoichiometryin subtropical China. Sci Rep. 2016; 6: 24261. DOI: 10.1038/srep24261

65. Griffiths PR, Madritch MD, Swanson AK. The effects of topography on forest soil characteristics in the Oregon Cascade Mountains (USA): Implications for the effects of climate change on soil properties. For Ecol Manag. 2009; 257(1): 1–7.

66. Bahn M, Rodeghiero M, Anderson-Dunn M, Dore S, Gimeno C, et al. Soil respiration in European grasslands in relation to climate and assimilate supply. Ecosystems. 2008; 11: 1352– 1367. DOI: 10.1007/s10021-008-9198-0

67. Shi Y, Baumann F, Ma Y, Song C, Uhn PK, Scholten T, He JS. Organic and inorganic carbon in the topsoil of the Mongolian and Tibetan grasslands: pattern, control and implications. Biogeosciences. 2012; 9: 2287–2299. doi:10.5194/bg-9-2287-2012

68. Vanhala P. Seasonal variation in the soil respiration rate in coniferous forest soils. Soil Biol Biochem. 2002; 34: 1375–1379.

69. Zhang YY, Wu W, Liu H. Factors affecting variations of soil pH in different horizons in hilly regions. PLoS One. 2019; 14(6): e0218563. doi: 10.1371/journal.pone.0218563

70. Martí C, Badía D. Characterization and classification of soils along two altitudinal transects in the Eastern Pyrenees, Spain. Arid Soil Res Rehabil. 1995; 9: 367–383. DOI: 10.1080/15324989509385905

71. Smith JL, Halvorson JJ, Jr HB. Soil properties and microbial activity across a 500 m elevation gradient in a semi-arid environment. Soil Biol Biochem. 2002; 34: 1749–1757.

72. Seibert J, Stendahl J, Sørensen R. Topographical influences on soil properties in boreal forests. Geoderma. 2007; 141(1–2): 139–148.

73. Badía D, Ruiz A, Girona A, et al. The influence of elevation on soil properties and forest litter in the Siliceous Moncayo Massif, SW Europe. J Mt Sci. 2016; 13: 2155–2169. https://doi.org/10.1007/s11629-015-3773-6

74. Klopfenstein ST, Hirmas DR, Johnson W. Relationships between soil organic carbon and precipitation along a climosequence in loess-derived soils of the Central Great Plains, USA. Catena. 2015; 133: 25–34. http://dx.doi.org/10.1016/j.catena.2015.04.015

75. Schrumpf M, Schulze ED, Kaiser K, Schumacher, J. How accurately can soil organic carbon stocks and stock changes be quantified by soil inventories?. Biogeosciences. 2011; 8: 1193–1212. https://doi.org/10.5194/bg-8-1193-2011.

76. Keenan TF, Migliavacca M, Papale D, Baldocchi D, Reichstein M, Torn M, Wutzler T. Widespread inhibition of daytime ecosystem respiration. Nat Ecol Evol. 2019; 3(3): 407–415. doi: 10.1038/s41559-019-0809-2

77. Reichstein M, Ciais P, Papale D, Valentini R, Running S, Viovy N. Reduction of ecosystem productivity and respiration during the European summer 2003 climate anomaly: a joint flux tower, remote sensing and modelling analysis. Glob Change Biol. 2007; 13(3): 634– 651. DOI 10.1111/j.1365-2486.2006.01224.x

78. Yu GR, Zhu XJ, Fu YL, He HL, Wang QF, Wen XF. Spatial patterns and climate drivers of carbon fluxes in terrestrial ecosystems of China Global Change Bio. 2013; l19: 798–810. https://doi.org/10.1111/gcb.12079

79. Chen S, Zou J, Hu Z, Lu Y. Climate and vegetation drivers of terrestrial carbon fluxes: a global data synthesis. Adv Atmos Sci. 2019; 36: 679–696.

80. Ma J, Liu R, Li C, Fan L, Xu G, Li Y. Herbaceous layer determines the relationship between soil respiration and photosynthesis in a shrub-dominated desert plant community. Plant Soil. 2020, 449: 193–207. https://doi.org/10.1007/s11104-020-04484-6

81. Raich JW, Tufekciogul A. Vegetation and soil respiration: Correlations and controls. Biogeochemistry. 2000; 48: 71–90. https://doi.org/10.1023/A:1006112000616

82. Chen Q, Wang Q, Han X, Wan S, Li L. Temporal and spatial variability and controls of soil respiration in a temperate steppe in northern China. Global Biogeochem Cycles. 2010; 24: GB2010. doi:10.1029/2009GB003538.

83. Oertel C, Matschullat J, Zurba K, Zimmermann F, Erasmi S. Greenhouse Gas Emissions From Soil - A review, Chemie der Erde. 2016; 76: 327–352, doi:10.1016/j.chemer.2016.04.002.

84. Andersson S, Nilsson SI. Influence of pH and temperature on microbial activity, substrate availability of soil-solution bacteria and leaching of dissolved organic carbon in a mor humus. Soil Biol Biochem. 2001; 33 (9): 1181–1191. https://doi.org/10.1016/S0038-0717(01)00022-0

85. Reth S, Reichstein M, Falge E. The effect of soil water content, soil Temperature, soil pH value and the root mass on soil CO2 effluxa modified model. Plant Soil, 2005; 268: 21–33.

86. Chappell C, Johnson A. Influence of pH and bulk density on carbon dioxide efflux in tree urban wetland types. Professional Agricultural Workers Journal. 2015; 3 (1, 5). http://tuspubs.tuskegee.edu/pawj/vol3/iss1/5

87. Mordhorst A, Peth S, Horn R. Influence of mechanical loading on static and dynamic CO2 efflux on differently textured and managed Luvisols. Geoderma. 2014; 219–220: 1–13.

88. Chen B, Liu S, Ge J, Chu J. Annual and seasonal variations of Q10 soil respiration in the sub-alpine forests of the Eastern Qinghai-Tibet Plateau, China. Soil Biol Biochem. 2010; 42: 1735–1742

89. Feng J, Wang J, Song Y, Zhu B. Patterns of soil respiration and its temperature sensitivity in grassland ecosystems across China, Biogeosciences. 2018; 15: 5329–5341, https://doi.org/10.5194/bg-15-5329-2018

90. Acosta M, Pavelka M, Montagnani L, Kutsch W, Lindroth A, Juszczak R, Janouš D. Soil surface CO2 efflux measurements in Norway spruce forests. Comparison between four different sites acrossEurope — from boreal to alpine forest. Geoderma. 2013; 192: 295–303. DOI: 10.1016/j.geoderma.2012.08.027.

91. Zhou T, Shi P, Hui D, Luo Y. Global pattern of temperature sensitivity of soil heterotrophic respiration (Q10) and its implications for carbon-climate feedback, J Geophys Res-Biogeo. 2009; 114: 271–274, https://doi.org/10.1029/2008JG000850, 2009.

92. Song X, Peng C, Zhao Z, Zhang Z, Guo B, Wang W, Jiang H, Zhu Q. Quantification of soil respiration in forest ecosystems across China. Atmos Environ. 2014; 94: 546–551, https://doi.org/10.1016/j.atmosenv.2014.05.071

93. Zhou W, Hui D, Shen W. Effects of Soil Moisture on the Temperature Sensitivity of Soil Heterotrophic Respiration: A Laboratory Incubation Study. PLoS ONE. 2014; 9(3): e92531. doi:10.1371/journal.pone.0092531

94. Zhao J, Li R, Li X, Tian L. Environmental controls on soil respiration in alpine meadow along a large altitudinal gradient on the central Tibetan Plateau. Catena. 2017; 159: 84–92. https://doi.org/10.1016/j.catena.2017.08.007

95. Zhou Z, Guo C, Meng H. Temperature sensitivity and basal rate of soil respiration and their determinants in temperate forests of North China. PLoS One. 2013; 8(12): e81793 https://doi.org/10.1371/journal.pone.0081793

96. Quan Q, Wang C, He N, Zhang Z, Wen X, Su H, Wang Q, Xue J. Forest type affects the coupled relationships of soil C and N mineralization in the temperate forests of northern China. Sci Rep. 2014; 4: 6584, DOI: 10.1038/srep06584,

97. Rambo TR, North MP. Canopy microclimate response to pattern and density of thinning in a Sierra Nevada forest. For Ecol Manag. 2009; 257 (2): 435–442, https://doi.org/10.1016/j.foreco.2008.09.029

98. Arx GV, Dobbertin M, Rebetez M. Spatio-temporal effects of forest canopy on understory microclimate in along-term experiment in Switzerland. Agr Forest Meteorol. 2012; 166–167: 144–155, https://doi.org/10.1016/j.agrformet.2012.07.018

99. Niu S, Luo Y, Fei S, Yuan W, Schimel D, Law BE, et al. Thermal Optimality of Net Ecosystem Exchange of Carbon Dioxide and Underlying Mechanisms. New Phytol. 2012; 194 (3): 775–783, DOI: 10.1111/j.1469-8137.2012.04095.x.

